# Measuring the non-selective effects of motor inhibition using isometric force recordings

**DOI:** 10.1101/2022.11.17.516968

**Authors:** Benjamin O. Rangel, Giacomo Novembre, Jan R. Wessel

## Abstract

Inhibition is a key cognitive control mechanism. When rapidly exerted, inhibitory control has broad, non-selective motor effects, typically demonstrated using cortico-spinal excitability measurements (CSE) elicited by transcranial magnetic stimulation (TMS). For example, during rapid action-stopping, CSE is suppressed at both stopped and task-unrelated muscles. While such TMS-based CSE measurements provide crucial insights into the fronto-basal ganglia circuitry underlying non-selective inhibition, they have several downsides. TMS is contraindicated in many populations (e.g., epilepsy, deep-brain stimulation patients), has limited temporal resolution, produces distracting auditory and haptic stimulation, is difficult to combine with other imaging methods, and necessitates expensive, immobile equipment. Here, we attempted to measure the non-selective motor effects of inhibitory control using a method unaffected by these shortcomings. 30 participants exerted isometric force on a hand-held force transducer while performing a foot-response stop-signal task. Indeed, when foot movements were stopped, force output at the task-irrelevant hand was suppressed as well. Moreover, this non-selective reduction of isometric force was highly correlated with stop-signal performance and showed frequency dynamics similar to established inhibitory signatures typically found in neural and muscle recordings. Together, we demonstrate that isometric force recordings capture the non-selective effects of motor inhibition, enabling many applications that are impossible with TMS.

## Introduction

Being able to stop actions is almost as essential as being able to start them. Our ancestors would not have gotten far had they been unable to stop reaching into a bush of berries once they saw a venomous snake in the underbrush. Inhibitory control allows humans to suppress prepared or ongoing motor processes when necessary. The deployment of inhibitory control in response to novel or unexpected stimuli must be rapid, so that potential threats can be appropriately reacted to. Previous work has identified a fronto-basal ganglia (FBg) network that implements rapid motor inhibition following salient stimuli (Aron and Poldrack 2006; Erika-Florence et al. 2014; Wager et al. 2005; Wessel and Aron 2017). This network includes the right inferior-frontal cortex and the pre-supplementary motor area, which rapidly activate the subthalamic nucleus (STN) of the basal ganglia via a hyper-direct fiber pathway – ultimately resulting in the suppression of thalamo-cortical output (Chen et al. 2020; Miocinovic et al. 2018; Nambu et al. 2002; Wessel and Aron 2017). Notably, when the FBg network exerts inhibition via this subthalamic hyper-direct pathway, it has broad, non-selective effects on the motor system. This is typically demonstrated using measurements of cortico-spinal excitability (CSE). CSE can be measured using a combination of Transcranial Magnetic Stimulation (TMS) and electromyography, evoking a motor-evoked potential that scales with the net excitability of the cortico-spinal tracts underlying specific muscle representations (Bestmann and Duque 2016; Lazzaro and Rothwell 2014; Volz et al. 2015). When humans exert inhibition to stop a specific action, CSE of the stopped muscle is indeed suppressed (Coxon et al. 2006; Leocani et al. 2000). However, most notably, this CSE suppression extends even to task-unrelated effectors (Badry et al. 2009). This non-selective effect of inhibitory control on CSE has been demonstrated using many different effector combinations (Cai et al. 2012; Wessel et al. 2013; Wessel and Aron 2017) and is causally attributable to the STN (Wessel et al. 2022).

Consequently, measurements of these non-selective effects of inhibitory control on CSE have been used to demonstrate the involvement of the fronto-subthalamic hyper-direct pathway in many different control-demanding scenarios, including those that do not explicitly require action-stopping. For example, unexpected perceptual stimuli (Dutra et al. 2018; Tatz et al. 2021), action errors (Guan and Wessel 2022), and response conflict (Wessel et al. 2019) have all been shown to be followed by short-latency, non-selective suppression of CSE – in line with prior work showing that all of these events activate the STN (Cavanagh et al. 2014; Frank et al. 2007; Herz et al. 2017; Siegert et al. 2014; Wessel et al. 2016). Finally, the assessment of CSE using TMS and EMG has been a popular and widespread method for studying not only inhibitory control, but also response preparation (Bestmann and Duque 2016; Leocani et al. 2000; Raud et al. 2020), interhemispheric interactions (Fiori et al. 2017; Hamada et al. 2014; Hannah and Rothwell 2017), and pathological states that affect the motor system (Badawy et al. 2013; Chowdhury et al. 2018; Jahanshahi and Rothwell 2017; Smith and Stinear 2016).

However, despite being an effective method for quantifying CSE and measuring the non-selective effects of inhibitory control on the motor system, the popular TMS-based CSE method has several substantial downsides. First, TMS is contraindicated in several populations, including, importantly, patients with a family history of epilepsy or other seizure disorders, those with implanted neurostimulators, or those with metal implants in their skull (Rossi et al. 2020). Many of these populations are most informative in regards to the associated fronto-basal ganglia brain networks underpinning inhibitory control, as human intracranial recordings are limited to exactly those patient groups. The contraindication of TMS in these groups is severely limiting inhibitory control research. Second, TMS-based CSE estimates have very poor temporal resolution, both due to the necessary recharging of monophasic TMS pulse generators (which rate-limits the collection of CSE measurements to around 1 Hz at most, Hallett and Chokroverty 2006), as well as the short- and long-term habituation effects of repeated TMS on CSE itself (Fitzgerald et al. 2006). Third, TMS pulses introduce unwanted tactile and auditory stimuli into an experiment (Rossi et al. 2020), which can influence behavior in many cognitive tasks. Fourth, the strong magnetic field produced by TMS can make it difficult to combine with other imaging methods. For example, attempts to combine TMS with electrocorticography (EEG) have proven successful, but the large artifacts introduced by the TMS pulses, as well as the subtle movements of the experimenter and/or participant in relation to the coil, can prove difficult to remove (Ilmoniemi et al. 2015; Rogasch et al. 2017). Finally, TMS stimulators are expensive and space-consuming, making them unfeasible to use in settings where mobility is key.

Therefore, we here aimed to evaluate a potential alternative method to quantify the non-selective effects of inhibitory control on the motor system – ideally, one that addresses these shortcomings of TMS-based CSE measurements. Specifically, we investigated whether non-selective inhibitory effects on the motor system could be found in the modulation of isometric force. This was motivated by several recent observations. First, previous research has shown that unexpected perceptual events trigger a multiphasic stimulus-locked modulation of isometric force (Novembre et al. 2018, 2019, Somerveil et al., 2021) – notably including a short-latency force reduction that is reminiscent of the fast and non-selective CSE suppression observed after such events (Dutra et al. 2018; Tatz et al. 2021). Importantly, this complex modulation of force is coupled with accompanying modulations of EEG activity, including a fronto-central positivity (Novembre et al. 2018), which has also been proposed as potential scalp-marker of the activity of the inhibitory FBg network (De Jong et al. 1990; Kok et al. 2004; Wessel and Aron 2015). Second, unexpected perceptual events also trigger a modulation of beta-like oscillatory activity (∼20 Hz) in the isometric force, which is coupled with isofrequent EEG activity (Novembre et al. 2019). Again, stopping-related activity in this particular frequency band is also found following stop signals (Swann et al. 2009; Wagner et al. 2018; Wessel 2020). Taken together, these previous observations led to the hypothesis that isometric force measurements may show the same non-selective suppression during action-stopping that is observed in CSE.

To test this hypothesis, we had 30 healthy adult participants perform a version of the Stop-Signal Task (SST; (Logan et al. 1984) in which responses were made via foot pedals. This same task had been shown to produce a clear pattern of non-selective CSE suppression at the task-unrelated hand in our prior TMS work (Tatz et al. 2021). However, instead of measuring CSE using TMS, participants exerted steady isometric pressure on a force transducer using the fingers of their task-unrelated right hand, while performing the stop-signal task with their feet. Our main hypothesis was that the isometric force output to these finger muscles would be transiently reduced when participants successfully withheld their foot responses, indicating the presence of non-selective inhibition. After indeed finding this pattern of results, we also correlated the magnitude of this effect to behavioral markers of stopping – specifically, to stop-signal reaction time (Verbruggen et al. 2019). Finally, we explored both the frequency-dynamics of the isometric force trace (Novembre et al. 2019) and its coefficient of variation, which has been associated with neuropsychiatric symptom severity in several clinical conditions (Davis et al. 2020; Hyngstrom et al. 2014), in relation to stopping behavior.

## Methods

### Participants

33 adult human participants were recruited for this study. Two datasets were incomplete due to hardware errors, and one additional participants’ data were rejected for violation of the race model (see behavioral analysis). This led to a final sample of 30 participants (age mean ± SD = 18.8 ± 1.6, 19 females, 29 right-handed). Participants were paid $15 per hour or received course credit for their participation in the study. All participants had normal or corrected-to-normal vision. The experiment was approved by the ethics committee at the University of Iowa (IRB #201511709).

### Experimental task and procedure

Stimuli were presented via an Ubuntu Linux computer, running Psychtoolbox 3 (Brainard 1997) under MATLAB 2017a (The MathWorks, Natick, MA). Participants sat upright with their arms resting on a supportive platform placed on the arm rests of the chair. Participants responded to stimuli during the task using left and right foot pedals (Kinesis Savant Elite 2). Participants performed a version of the classic stop-signal task (Logan et al. 1984), while attempting to maintain a constant pinch-grip force with their right hand (*Figure 1*). Stop-signal task stimuli were presented on an all-grey background. All trials began with a black fixation cross (500ms), followed by a black arrow (1000ms) pointing left or right (Go-Signal). Participants were instructed to respond to the Go-Signal using foot pedals (left pedal for left facing arrows, and right pedal for right facing arrows) using both feet, within the 1000ms that the arrow was displayed. The arrow would immediately disappear following a response. The Go-Signal was followed by a variable length intertrial interval (ITI) which ensured all trials had a total length of 3s. If participants responded to the Go-Signal, a black fixation cross would be displayed during the ITI, if they did not respond, the message “TOO SLOW” was displayed in red text instead. During 1/3 of trials, the Go-Signal was followed by a Stop-Signal (i.e., the black arrow turning red) after a stop-signal delay (SSD). Participants were instructed to withhold their response on these trials. The SSD began at 200ms and was subsequently adjusted in steps of 50ms (added to the SSD after successful stop trials and subtracted after failed stop trials) with the goal of participants successfully stopping their responses on half of all stop-trials (Verbruggen et al. 2019). Participants performed 10 blocks, each of 60 trials. At the end of each block, performance feedback was displayed on the screen. Participants were instructed that responding quickly on Go-trials and stopping successfully on Stop-trials were equally important.

**Figure 1.**
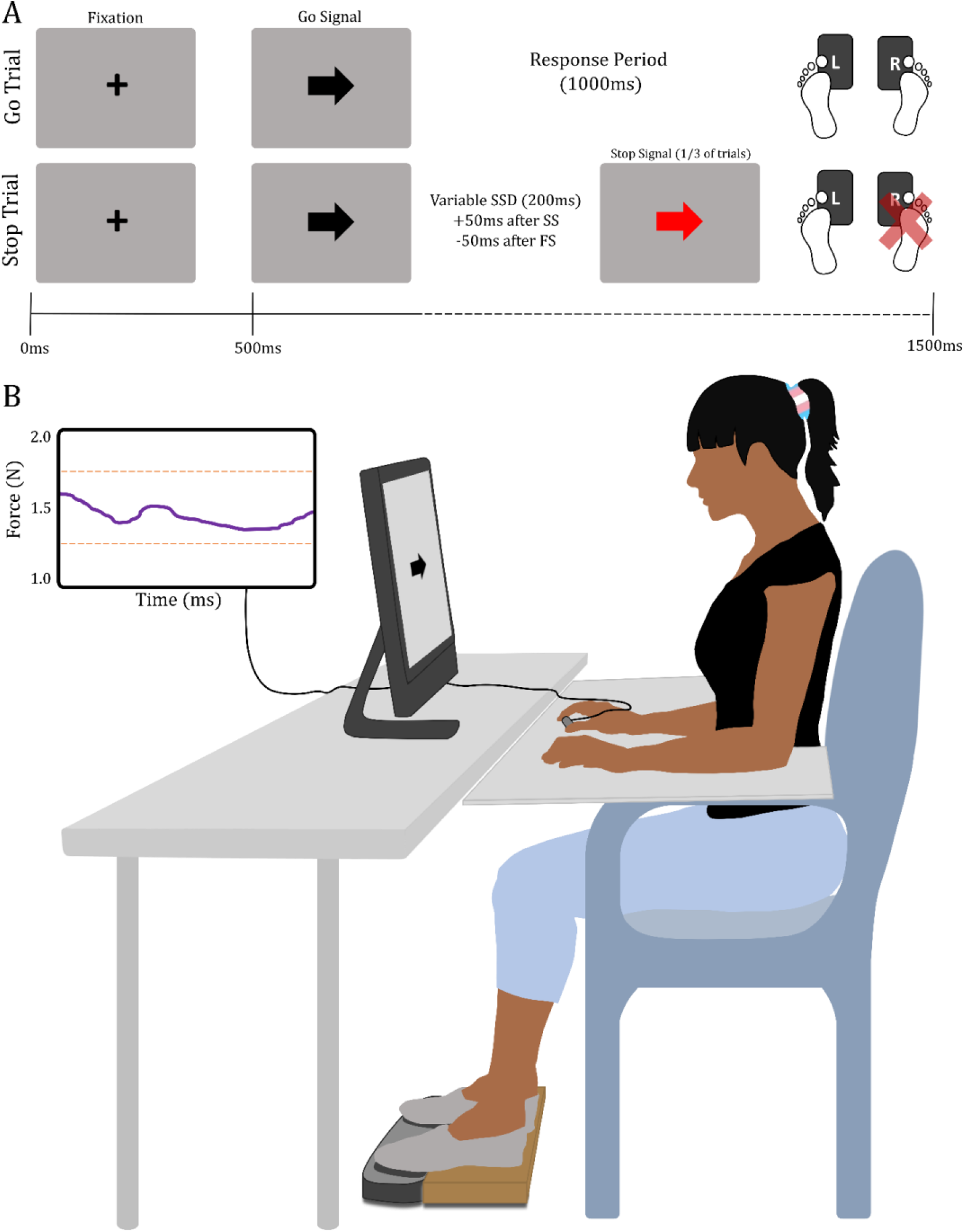
Paradigm and task diagram. A) During both Stop and Go trials (2/3 of trials), the fixation cross was presented for 500ms, before the onset of the Go signal (black arrow). Subjects then had 1000ms to make a response, using the foot pedals. During Stop trials, the Go signal was followed by the Stop signal (red arrow) following a variable SSD, beginning at 200ms (+50ms after successful stop, -50ms after failed stop). B) All subjects held the force transducer in their right hand, between their index finger and thumb. Their arms were supported on a wooden platform, which rested on the chair arms. A wooden board was placed under the subject’s feet to elevate their toes above the foot pedals and prevent fatigue. Live force recordings were presented on a second computer screen, which was outside of the subjects’ direct view. Prior to each block, subjects would direct their view towards the second screen to recenter their force output ∼1.5N.

### Force Recording

Participants held the force transducer between their right index finger and thumb throughout the blocks of the task. Live force data was recorded using a highly sensitive force/torque sensor (Nano17, ATI Industrial Automation), with custom 3D printed finger grips attached to the sensor. The live force data were recorded and displayed via a separate Windows 7 computer running MATLAB 2020b, using custom scripts. Force was recorded at a rate of 3000Hz and then resampled to a rate of 1000Hz for display purposes. The monitor showing the force data was placed on a shelf out of direct view of the participants, except for the calibration time periods between blocks. A twist-tie secured the bottom transducer grip to their thumb, and they rested their forearm and hand on the supportive platform placed on the arm rests of the chair. Prior to the beginning of each task block, participants were instructed to direct their gaze towards the second screen displaying the live force readings. They were instructed to stabilize their force output within the range of 1.25-1.75N (*Figure 1b*). Once they felt that they were steady enough, the participant would look back towards the task screen, and the experimenter would start the task block. Participants wore a sun visor to obscure any peripheral view of the second screen when performing the task.

### Behavioral Analysis

Behavioral data were processed following consensus protocols for analysis of the stop-signal task (Verbruggen et al. 2019). Trials with incorrect responses, missing responses on Go trials, responses faster than 150ms, or where the foot pedal was prematurely depressed, were rejected (0.64% of trials [0 — 2.5%]). Blocks where the proportion of successful stops fell below.25, or exceeded.75, were also rejected (this was the case for 1 block out of 10 in n=3, 2 blocks in n=1, and 3 blocks in n=1). Participants’ stop-signal reaction time (SSRT) was then calculated using the integration method (Matzke et al. 2018).

### Force Data Analysis

Single subject raw data were bandpass filtered (0.1 – 40Hz) before being epoched relative to the arrow onset (-500 – 2500ms). Any trial rejected during the behavioral analysis was removed from the force analysis as well. The remaining trials were baseline corrected to the -250-0ms pre-arrow period. Trial epochs were screened for outliers within the -250 to 850ms time window. Specifically, trials with large fluctuations in force (outside -0.3 – 0.3N) were removed (0.54% of trials [0 – 3%]. Then, the remaining trials were screened for force fluctuations outside 4 SDs of the mean of all trials, regardless of condition (4.0% of trials [2 – 6.2%], cf., Novembre et al., 2019). The mean percentage of trials rejected for subjects was 8.0% [2 – 19.3%] for subjects without block rejections. The rejection percentages are listed in *Table 1*. Participant data were then averaged by condition. In a separate analysis, we epoched force data relative to the response (-1500 – 1500ms). These trials were baseline corrected to the -1500ms to - 750ms pre-response period. Further processing was done as above.

**Table 1.** Mean (SD) and range [min max] of the percent of trials rejected per criteria. ‘Pedal down’ criterion represents trials where the pedal was depressed prior to stimulus presentation. Shaded areas denote the average percent of trials rejected by behavioral and force criteria. Totals were also calculated for those who did and did not have blocks rejected due to the proportion of successful stops falling below.25, or exceeding.75.

|  | % mean (SD) |  | % range |  |
| --- | --- | --- | --- | --- |
| No Response | 2.0 | (2.1) | [0 | 8.8] |
| Response Error | 0.5 | (0.5) | [0 | 2.0] |
| Fast RT (< 150ms) | 0.3 | (0.4) | [0 | 1.5] |
| Pedal Down | 0.5 | (0.8) | [0 | 3.3] |
| Force outside $\pm 0.3N$ | 0.5 | (0.7) | [0 | 3.0] |
| Force outside 4*SD | 4.0 | (1.1) | [2 | 6.2] |
| Total behavioral (all blocks, n = 25) | 3.5 | (2.8) | [0 | 12.2] |
| Total behavioral (blocks rejected, n = 5) | 18.4 | (8.9) | [11 | 32.2] |
| Total force | 4.6 | (1.6) | [2 | 7.5] |

### Time-Frequency Analysis of the force trace

Filtered, continuous force data were convolved with a complex Morlet wavelet (3-10 cycles), for 40 linearly spaced frequencies (1-30Hz; Cohen 2014). Power was estimated using the squared magnitude of the complex wavelet-convolved data. Time-frequency power estimates were then epoched relative to the arrow and response (same time span as in the force data). Trials marked as behavioral rejections, and outliers identified during the force data analysis, were removed. Baseline correction (decibel conversion) were performed per condition for arrow-locked (-200-0ms) and response-locked (-1500 to -750ms) power estimates, before being averaged.

### Beta-Burst Analysis

Raw force data were passband filtered (0.3 – 30) prior to being convolved with a complex Morlet wavelet (7 cycles), for 15 linearly spaced frequencies (15-29Hz) in the beta range (Wessel, 2020). Note that this decomposition differs slightly from the one above to match established methods in the beta burst literature. The squared magnitude of the complex wavelet-convolved data was used to estimate the power. The estimates were then epoched relative to the arrow and response (same time spans as in the force data). Trials marked as behavioral rejections, and outliers identified during the force data analysis, were removed. Individual peaks in beta power were detected during single trials, using the *imregionalmax* Matlab function. A peak in beta power was counted as a burst if its amplitude exceeded 3X the median power for that frequency across all trials, based on methods used in (Bräcklein et al. 2022).

### Coefficient of the Variation of Force (CVF)

Unfiltered force data were epoched relative to the arrow onset (-500 – 2500ms). Behavioral rejections and trials rejected during the force analysis were removed. CVF was calculated for each trial, across all time points (*σ*/*μ* * 100), producing a single CVF value per trial. CVF values were then averaged across trials, to yield a single CVF value per subject, as measure of individual force unsteadiness. These values were then correlated with subjects’ SSRT (Pearson’s). A single subject was rejected as an outlier based on Cook’s distance (Cook 1977), leading to the final correlation containing 29 subjects.

## Results

### Behavioral Results

The reaction time (RT) results are displayed in *Figure 2*. Subjects’ mean Go RT (580ms), failed-stop RT (513ms), and SSRT (309ms) all showed significant differences from each other when compared using paired-samples t-tests (Go vs FS: t(29) = 19.9, p <.001; Go vs SS: t(29) = 16.7, p <.001; FS vs SS: t(29) = 13.3, p <.001).

**Figure 2.**
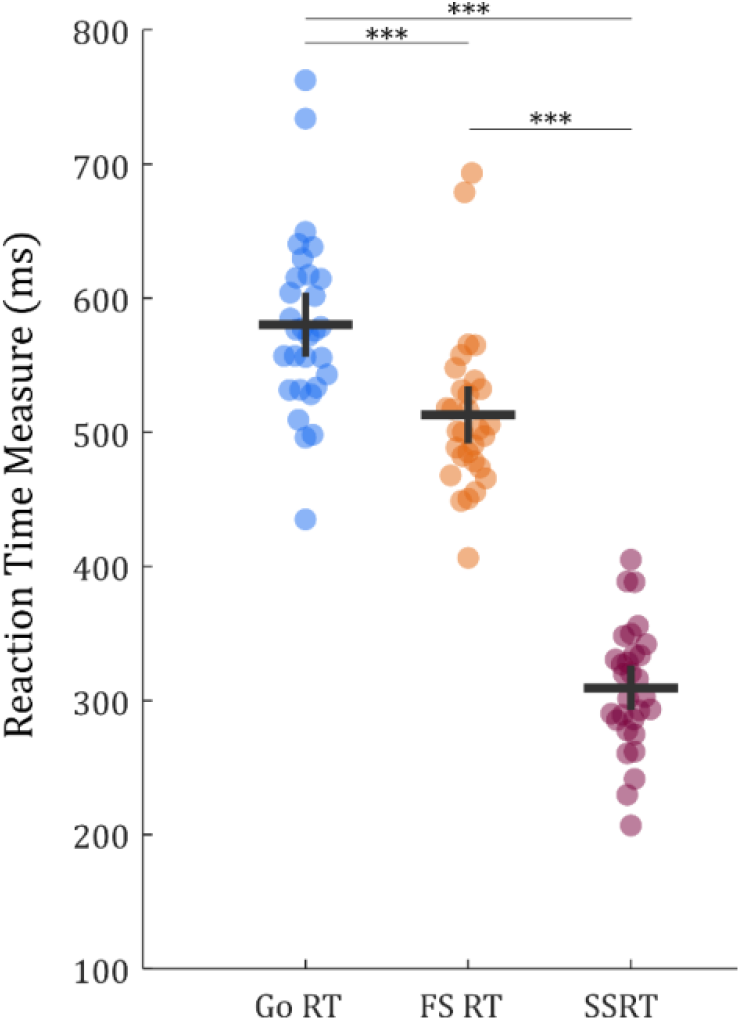
Colored dots denote single subjects’ RT measures. Horizontal and vertical lines represent mean and SEM, respectfully. All three conditions differed from each other significantly (*** p <.001).

### Isometric force

Significant differences in force output were observed between the three conditions of interest in the following time-ranges, corrected for multiple comparisons using the FDR method (*Figure 3a*): *Go vs. successful Stop (31-64ms, 183-390ms, 411-643ms, 1350-2500ms), Go vs. failed Stop (1-2500ms) and failed vs. successful stop (1-2079ms*). Qualitatively, the force trace for successful Stop-trials (SS) showed a triphasic pattern highly similar to previous findings where unexpected stimuli were presented to participants during isometric force exertion (Novembre et al. 2018). Here, the pattern was slightly time-delayed, likely due to the visual nature of the stop-signal (compared to the lower latency auditory and haptic stimuli used in Novembre et al., 2018). Notably, the initial dip of this pattern (∼150-250ms post-arrow) – which was larger (more negative) for successful Stop-trials compared to both Go- and failed Stop-trials (FS) and was largely absent in the force traces of Go and failed Stop trials – preceded the mean SSRT (309ms post-arrow) of all subjects.

**Figure 3.**
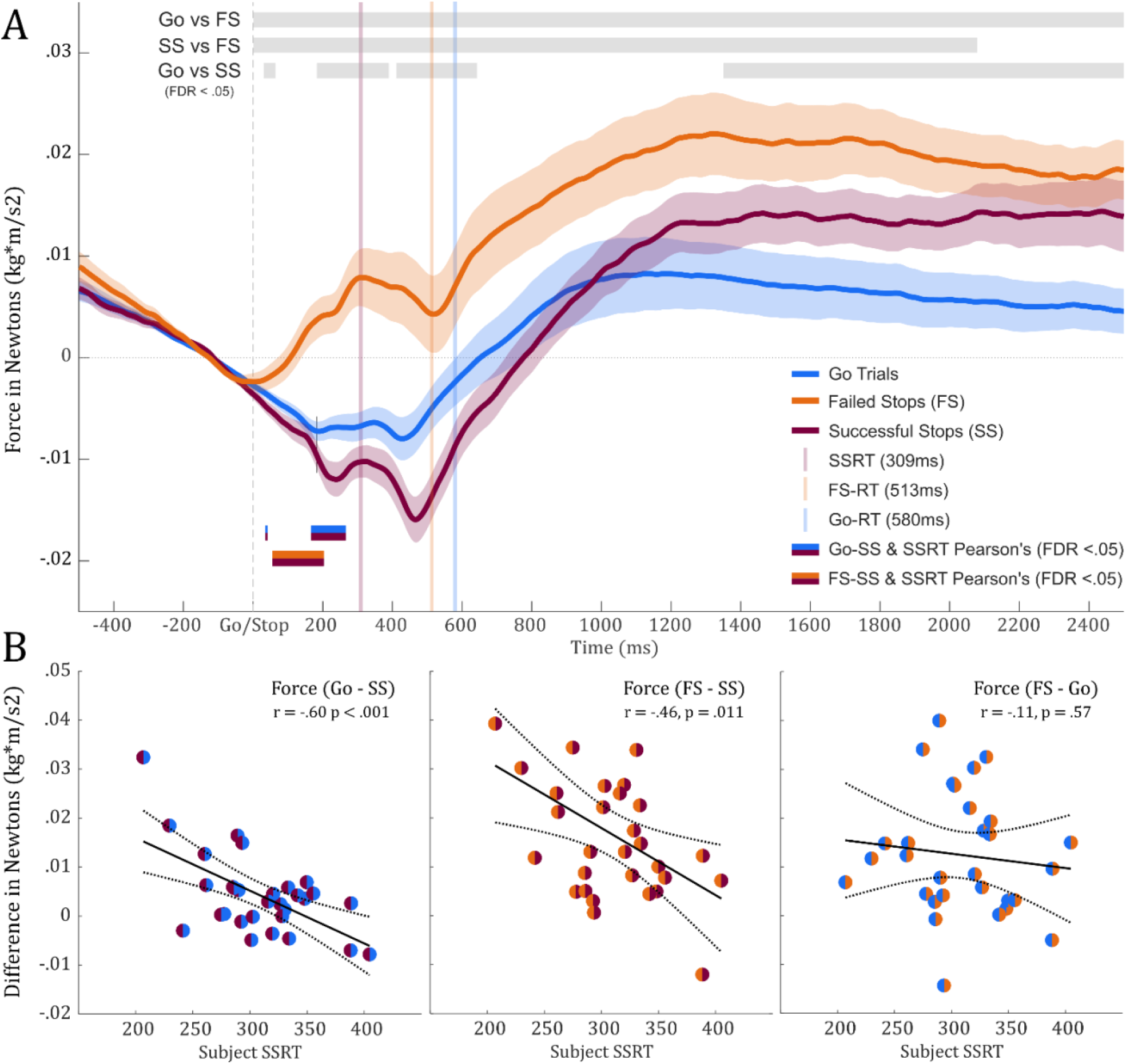
A) Plot shows the grand mean of force traces for all three conditions (Go, FS, SS), time-locked to the presentation of the stimulus. Significant differences (FDR <.05) between traces are denoted by grey bars at the top of the plot. Double-colored bars at the bottom of the plot indicate when individual differences in force between conditions significantly correlate with single subject SSRT. B) Pearson’s correlations of the differences between conditions, from 183ms post-arrow (highlighted by the vertical line intersecting the Go and SS traces) to mean SSRT (309ms post-arrow) for each subject, and subjects’ SSRT.

Pearson correlations between the force differences of the conditions (Go-SS, FS-SS, FS-Go) and subjects’ SSRT were calculated at every timepoint, with FDR-corrected significant periods highlighted in *Figure 3a* (*colored bars;* time-ranges: Go-SS [36-41ms, 167-266ms] and FS-SS [55-503ms]). To generate a scatter plot, the average differences between conditions were calculated from 183ms post-arrow (where the initial dip on successful Stop trials began to significantly diverge from the Go condition) to 309ms post-arrow (the mean SSRT in this sample) and then correlated to SSRT (*Figure 3b*). SSRT was highly correlated with differences in force output between Go/SS trials (r = -.61, p <.001) and FS/SS trials (r = -.48, p =.006), during this time period.

The unexpected morphology of the failed Stop-trial trace – especially its early divergence from the successful Stop and Go-trials – suggests that pre-signal differences may distinguish those trials from the other two conditions. *Supplementary Figure 1* (Go-trial locked force traces for all three conditions) shows that this is not the case, as Go- and Failed-Stop trials looked highly similar. Moreover, while the Stop-locked activity was of primary a priori interest in this study, the Go-locked data showed the same force reduction in the successful Stop-condition (with additional smearing due to the varying Stop-signal delay blurring the triphasic pattern observed in the Stop-signal locked data, making the effect look smoother, cf., *Supplementary Figure 1*).

### Time-frequency analysis of force trace

Following time-frequency decomposition of the force timeseries, we observed stimulus-induced modulations of specral power in both low (∼4Hz – Go trials) and high (13-30 Hz – all trials) frequencies, consistently with the results reported by (Novembre et al. 2019). Notably, distinct patterns of spectral power can be observed between Go and SS trials. SS trials showed the greatest increase beta-band power following stimulus presenation, but prior to the mean SSRT (*Figure 4a*). Contrasts between conditions showed that the significant differences between Go and SS trials, prior to SSRT, fall almost entirely within the beta-band range (mean f(Hz) = 17.4 ± 2.9; *Figure 4b*). Power differences between conditions were averaged across all time points prior to mean SSRT (309ms post-arrow) and frequencies within the beta-band range (13-30Hz) to produce a single value per contrast and per subject. These values were then correlated with subjects’ SSRT (Pearson’s; *Figure 4b*). Greater differences in power between SS and Go trials predicted faster SSRTs (r =.46, p =.011). Comparisons between successful Stop and failed Stop-trials can be found in *Supplementary Figure 2*.

**Figure 4.**
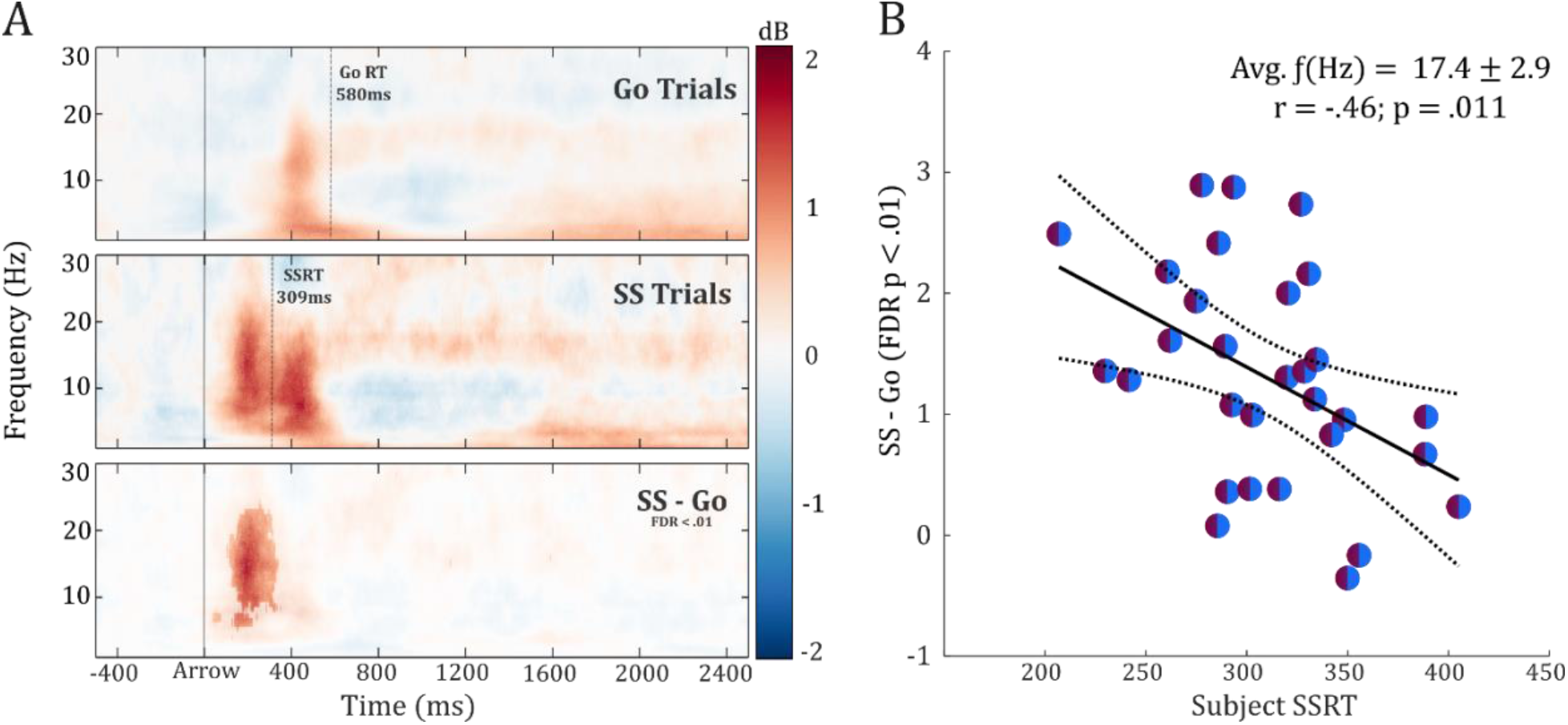
Results from the time-frequency analysis. A) Plots show the spectral power of the force trace, averaged for Go and SS conditions across subjects, for frequencies 1-30Hz. Areas of significant difference between Go and SS conditions (FDR <.01) are highlighted in the third panel. B) The differences between conditions were averaged across all post-arrow/pre-SSRT time-points and all frequencies within the β-band (13-30Hz), and compared to single subject SSRT, using Pearson’s correlations.

### β-Burst rate analysis of isometric force trace

β-burst rate differences were observed during the timespan of interest (0-600ms post-arrow, *Figure 5b*). A consistent increase in β-burst rate can be observed in SS trials, compared to Go trials, from 150-250ms, which temporally coincides with the significant relationship their force difference (Go-SS) and subject SSRT (*Figure 3a*). SS trials show a greater β-burst rate than FS trials, during a narrower timespan (150-200ms post-arrow), before the burst rate for FS trials approaches that of the SS trials in the 200-225ms time bin.

**Figure 5.**
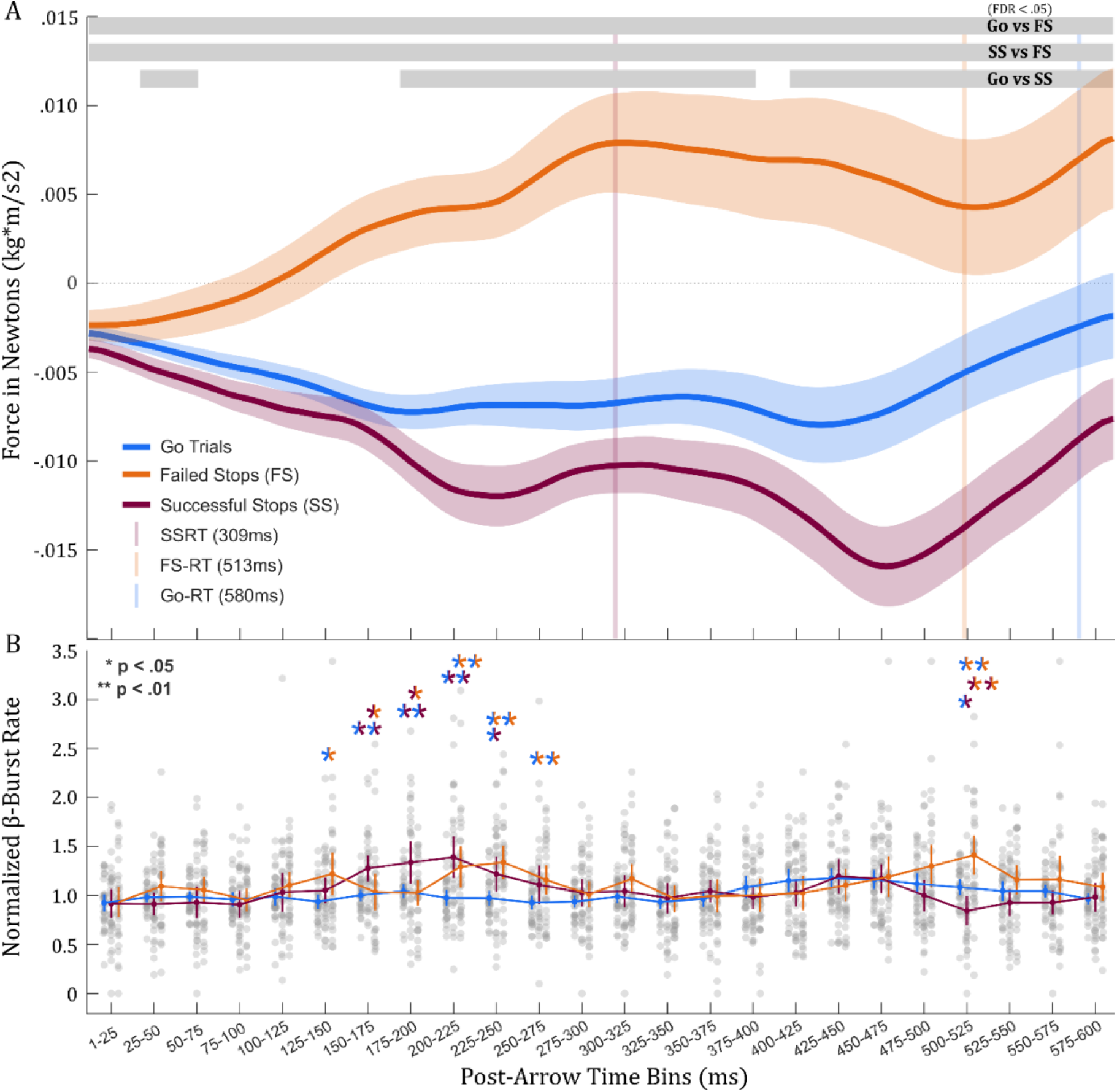
A) Plot is a zoomed in version of Figure 3a – i.e., the time-domain representation of the force trace, limited to the time-points 0-600ms post-arrow. Grey bars at the top of the plot indicate signifcant differences between condititons in the time domain (FDR <.05). B) Normalized β-burst rates of force data, averaged for each condition, across subjects. Semi-transparent grey dots denote individual subjects’ data points. Colored dots represent the mean within each time bin, and are intersected by vertical lines indicating the SEM.

### Coefficient of Variation of Force (CVF)

The mean coefficient of variation of force was computed across all time points (-500 pre-arrow to 2500ms post-arrow) and across all trials, for each subject. Following the rejection of one subject (cook’s =.71, *Figure 6b*), the remaining 29 subjects’ CVF and SSRTs were compared using a Pearson’s correlation. Greater motor steadiness (lower CVF) signifcantly predicted faster SSRTs (*Figure 6a*).

**Figure 6.**
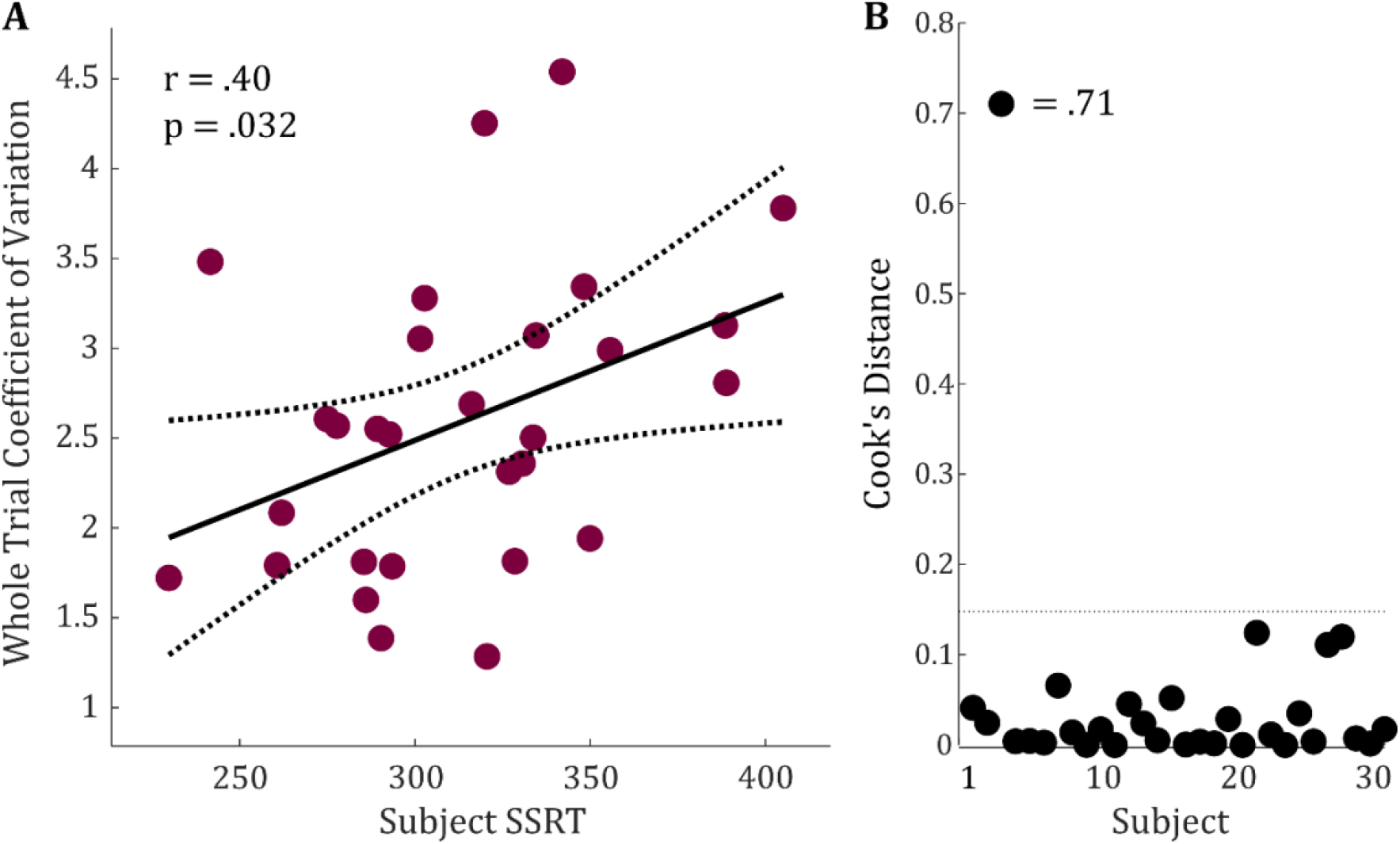
The relationship between the mean coefficient of variation of force of each subject and their SSRTs. A) The mean whole-trial (-500 to 2500ms post-arrow) CVF for each subject was correlated with their SSRTs (Pearson’s, r =.40, p =.032), after the rejection of one outlier (B) based on Cook’s Distance.

## Discussion

In the current study, we tested whether measurements of isometric force can be used to capture the non-selective effects of inhibitory control on the motor system. Indeed, in line with our prediction, data from a foot-response stop-signal task showed that successful Stop-trials yielded a significant force suppression at the task-unrelated hand prior to stop-signal reaction time, paralleling existing reports of non-selective CSE suppression during action-stopping. Moreover, the magnitude of this suppression was highly predictive of SSRT itself, with subjects that showed faster SSRT also showing stronger short-latency suppression of isometric force. Finally, the force data contained highly useful additional information. First, time-frequency decompositions of the force trace showed a preponderance of beta-band activity during stopping, in line with converging evidence for the importance of this frequency band in motor processes coming from other imaging domains (see below). Second, the coefficient of variation across the entire dataset was also correlated with SSRT. This has important clinical implications, as the CVF has been shown to be elevated in the elderly (Enoka et al. 2003; Galganski et al. 1993) and in Parkinson’s Disease (Skinner et al. 2019), as well as predictive of poorer coordination (Almuklass et al. 2016) and balance in several clinical populations (Davis et al. 2020; Hyngstrom et al. 2014).

Our main findings mirror those of previous work demonstrating CSE suppression during the stop-signal task. Previous research using TMS has shown that CSE is non-selectively suppressed when participants are able to successfully stop their responses (Badry et al. 2009; Tatz et al. 2021; Wessel et al. 2013; Wessel and Aron 2017). This suppression is observed approximately 150ms after the presentation of the stop-signal (Jana et al. 2020; Majid et al. 2012; Wessel and Aron 2017). We observed a commensurate decrease in force output during successful-stop trials in the current study. Indeed, this non-selective suppression of isometric force emerged almost exactly 150ms post-arrow, becoming statistically significant at 183ms (*Figure 3a*). Furthermore, the individual differences in force output between conditions during this initial suppression period were negatively correlated with single-subject SSRT, beginning at 167ms. Thus, isometric force recordings provide a temporally precise and highly accurate method for the assessment of non-selective motor suppression during inhibitory control. This is important because force recordings are not affected by many of the common shortcomings associated with TMS-based measurements of CSE. First, unlike TMS methods, force recordings are continuous and allow a quantification of non-selective suppressive effects with dense temporal coverage. This is particularly important since there are meaningful differences in the latency with which the underlying inhibitory control system operates (Chen et al. 2020; Coxon et al. 2012). TMS only allows the collection of a single sample of CSE at one specific time point per trial. Hence, different subjects may show maximal suppression of CSE at slightly different time points. Force recordings can detect such variations by continuously sampling throughout the trial. Second, unlike TMS-based methods, force recordings do not introduce auditory and haptic stimulation to the subject, and are hence potentially less distracting. Third, force recordings do not introduce stimulation artifacts into concurrent neural recordings, and hence do not interfere with other simultaneously acquired data. Fourth, force recordings can be readily performed in populations in which TMS is contraindicated, for example, those with epilepsy, a family history of seizure disorders, or implanted medical devices. Crucially, this can also enable investigations of non-selective motor inhibition using imaging techniques that typically depend on such populations (such as ECoG, which is only done in epilepsy patients). Fifth, force recordings are comparatively cheap and can be done with minimal footprint. As such, they present a highly viable alternative to TMS-based investigations of the non-selective effects of motor inhibition. Of course, TMS-based methods have other advantages – primarily, the fact that the underlying neuronal dynamics of CSE (and other TMS-based indices like intracortical inhibition) are relatively well understood. Future usage of force or TMS-based methods for the quantification of non-selective inhibition will depend on the exact research question, and future research should investigate their potential correspondence.

Another advantage of recording densely sampled time-series data from a force transducer is the ability to explore the dynamics of these data in the frequency domain (Novembre et al., 2019). Cortical oscillations in the β-band (13-30Hz), have long been associated with motor function. A prominent desynchronization of β-band power is observed during movement initiation (McFarland et al. 2000; Pfurtscheller et al. 2003), while increases in β-band power are observed during the cancellation of movements (Picazio et al. 2014; Soh et al. 2021; Swann et al. 2012; Wagner et al. 2018; Wessel 2020). Here, we found that significant differences between conditions by decomposing isometric force data into the time-frequency domain. Most notably, we found that successful Stop-trials show increased β-band activity compared to Go-trials, and that these differences predict single-subject SSRT (*Figure 4*). Recently, the measurement of cortical β-band activity has shifted towards single-trial estimations of burst-like events, rather than trial-averaged changes in power. Previous research has shown that β-activity is characterized as transient bursts, rather than prolonged changes (Bonaiuto et al. 2021; Feingold et al. 2015; Leventhal et al. 2012; Sherman et al. 2016), and that the presence or absence of these β-bursts better predicts behavior (Shin et al. 2017; Soh et al. 2021; Wessel 2020). We compared the rate of β-bursts for each condition, and showed that the bursting rate for successful-stop trials was significantly greater than those of failed Stop- and Go trials, from 150-200ms post-arrow (*Figure 5*). A delayed increase in β-burst rate for failed Stop- trials followed by about 50ms. This finding is consistent with the idea that burst rates indicate more successful movement cancelation.

Finally, our exploratory analysis of the coefficient of variation of force (CVF) highlights additional potential applications of isometric force measurements. CVF of isometric force data has long been associated with a variety of clinical measures (Enoka and Farina 2021). Greater CVF has been shown to accompany poorer performance on walking tests (Mani et al. 2018), grooved pegboard tests (Almuklass et al. 2016; Feeney et al. 2018), and assessments of postural sway (Kouzaki and Shinohara 2010), especially in elderly patients. Increased CVF has also been associated with greater symptomology in Parkinson’s disease (Wilson et al. 2020), stroke survivors (Hyngstrom et al. 2014) and patients with multiple sclerosis (Davis et al. 2020). While some studies have explored the modulation of CVF due to task conditions or stimuli presentation (Christou et al. 2004; Farina et al. 2012), it remains relatively unexplored in relation to higher cognition and non-clinical assessments. Here, we demonstrated that the averaged whole-trial CVF for each subject was positively correlated with SSRT. Hence, performance metrics in the stop-signal task may provide a potential complementary window into these clinical assessments.

In sum, we here report a novel signature of the non-selective effects of inhibitory control on the motor system. Isometric force recordings from a task-unrelated motor effector showed a clear reduction in force when another effector was successfully stopped. Moreover, the degree of this suppression was highly correlated with participants’ stopping ability. Unlike the previous gold-standard method used to demonstrate such non-selective motor effects (TMS-based CSE recordings), isometric force recordings provide high temporal resolution, superior compatibility with other imaging methods, applicability in previously inaccessible populations, minimal distraction to the subject, a small footprint, and low cost. Future work should capitalize on these properties to study the neural underpinnings of the non-selective effects of inhibitory control on the motor system.

## Acknowledgements

This work was funded by the National Science Foundation (NSF CAREER 1752355 to JRW).

## Supplementary Materials

**Supplementary Figure 1.**
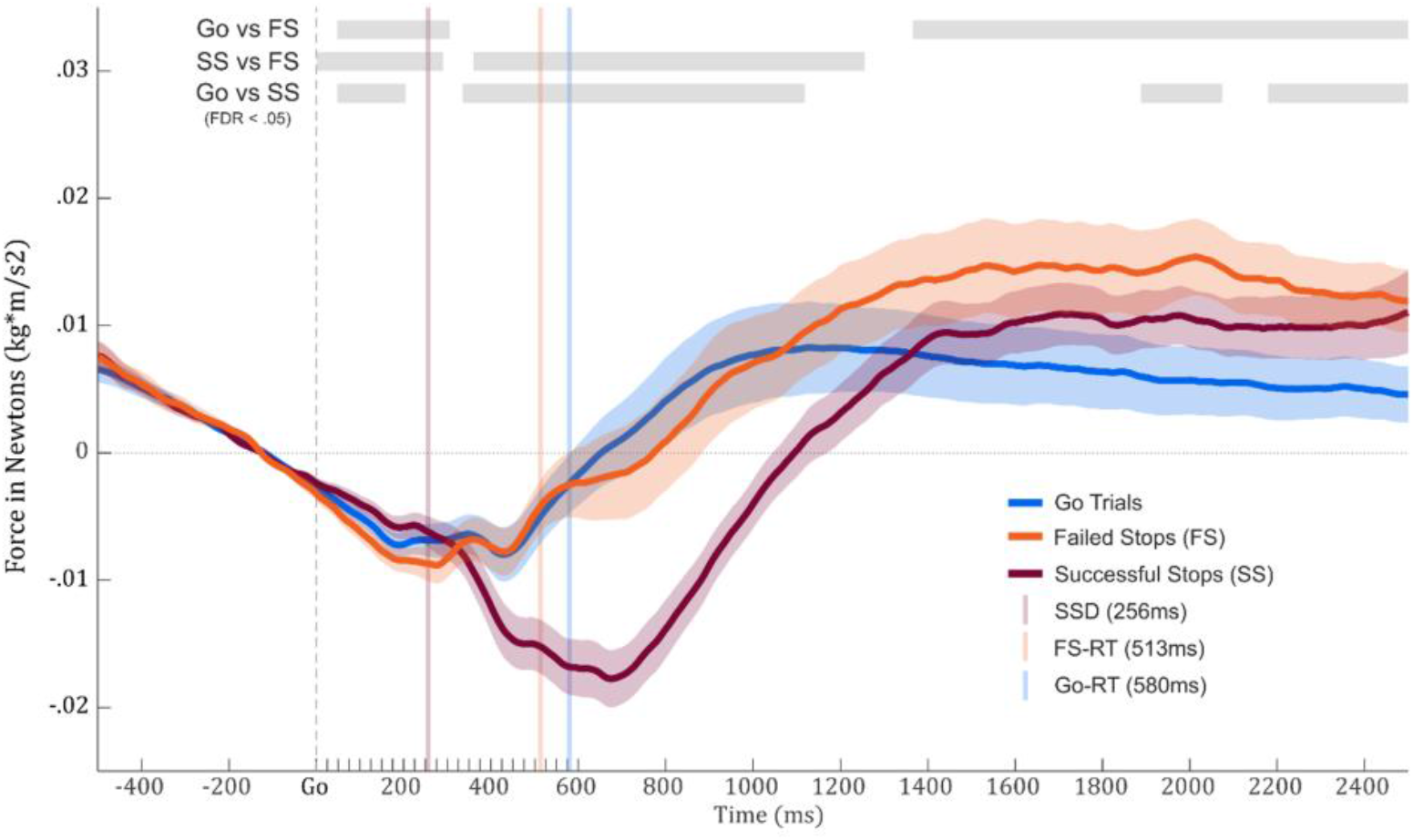
Isometric force traces time-locked to the Go-signal

**Supplementary Figure 2.**
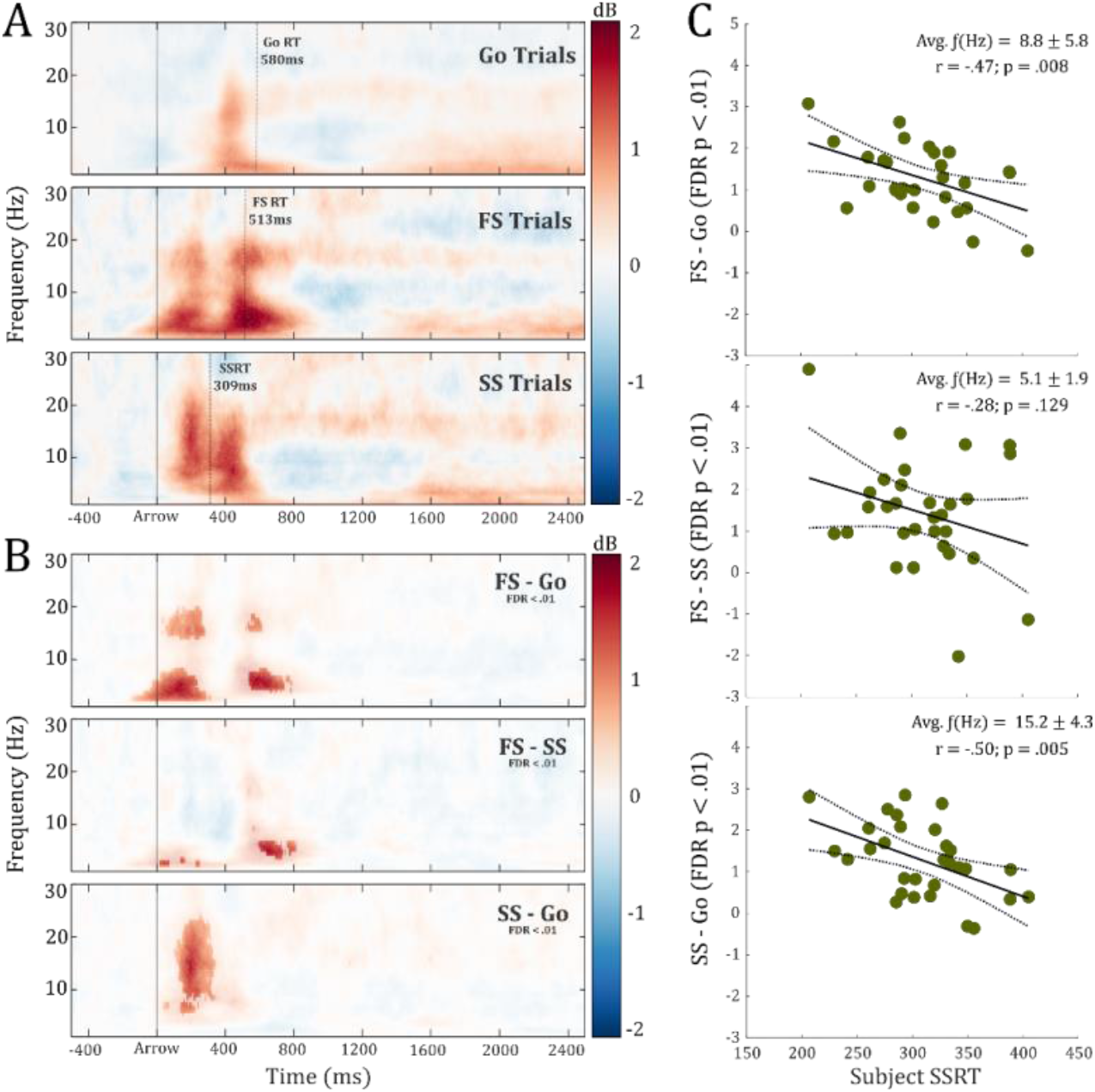
Time-frequency comparisons for successful Stop, Go- and failed Stop-trials.

## References

Almuklass AM, Price RC, Gould JR, Enoka RM. Force steadiness as a predictor of time to complete a pegboard test of dexterity in young men and women. J Appl Physiol 120: 1410–1417, 2016.

Aron AR, Poldrack RA. Cortical and Subcortical Contributions to Stop Signal Response Inhibition: Role of the Subthalamic Nucleus. J Neurosci 26: 2424–2433, 2006.

Badawy RAB, Jackson GD, Berkovic SF, MacDonell RAL. Cortical excitability and refractory epilepsy: a three-year longitudinal transcranial magnetic stimulation study. Int J Neural Syst 23: 1250030, 2013.

Badry R, Mima T, Aso T, Nakatsuka M, Abe M, Fathi D, Foly N, Nagiub H, Nagamine T, Fukuyama H. Suppression of human cortico-motoneuronal excitability during the Stop-signal task. Clin Neurophysiol 120: 1717–1723, 2009.

Bestmann S, Duque J. Transcranial Magnetic Stimulation: Decomposing the Processes Underlying Action Preparation. Neurosci 22: 392--405, 2016.

Bonaiuto JJ, Little S, Neymotin SA, Jones SR, Barnes GR, Bestmann S. Laminar dynamics of high amplitude beta bursts in human motor cortex. Neuroimage 242: 118479, 2021.

Bräcklein M, Barsakcioglu DY, Vecchio AD, Ibáñez J, Farina D. Reading and Modulating Cortical β Bursts from Motor Unit Spiking Activity. J Neurosci 42: 3611–3621, 2022.

Brainard DH. The Psychophysics Toolbox. Spatial Vision 10: 433–436, 1997.

Cai W, Oldenkamp CL, Aron AR. Stopping speech suppresses the task-irrelevant hand. Brain Lang 120: 412–415, 2012.

Cavanagh JF, Sanguinetti JL, Allen JJB, Sherman SJ, Frank MJ. The Subthalamic Nucleus Contributes to Post-error Slowing. J Cognitive Neurosci 26: 2637–2644, 2014.

Chen W, Hemptinne C de, Miller AM, Leibbrand M, Little SJ, Lim DA, Larson PS, Starr PA. Prefrontal-Subthalamic Hyperdirect Pathway Modulates Movement Inhibition in Humans. Neuron 106: 579-588.e3, 2020.

Chowdhury NS, Livesey EJ, Blaszczynski A, Harris JA. Variations in response control within at-risk gamblers and non-gambling controls explained by GABAergic inhibition in the motor cortex. Cortex 103: 153–163, 2018.

Christou EA, Jakobi JM, Critchlow A, Fleshner M, Enoka RM. The 1-to 2-Hz oscillations in muscle force are exacerbated by stress, especially in older adults. J Appl Physiol 97: 225–235, 2004.

Cook RD. Detection of Influential Observation in Linear Regression. Technometrics 19: 15–18, 1977.

Coxon JP, Impe AV, Wenderoth N, Swinnen SP. Aging and Inhibitory Control of Action: Cortico-Subthalamic Connection Strength Predicts Stopping Performance. J Neurosci 32: 8401–8412, 2012.

Coxon JP, Stinear CM, Byblow WD. Intracortical Inhibition During Volitional Inhibition of Prepared Action. J Neurophysiol 95: 3371–3383, 2006.

Davis LA, Alenazy MS, Almuklass AM, Feeney DF, Vieira T, Botter A, Enoka RM. Force control during submaximal isometric contractions is associated with walking performance in persons with multiple sclerosis. J Neurophysiol 123: 2191–2200, 2020.

Dutra IC, Waller DA, Wessel JR. Perceptual Surprise Improves Action Stopping by Nonselectively Suppressing Motor Activity via a Neural Mechanism for Motor Inhibition. J Neurosci 38: 1482–1492, 2018.

Enoka RM, Christou EA, Hunter SK, Kornatz KW, Semmler JG, Taylor AM, Tracy BL. Mechanisms that contribute to differences in motor performance between young and old adults. J Electromyogr Kines 13: 1–12, 2003.

Enoka RM, Farina D. Force Steadiness: From Motor Units to Voluntary Actions. Physiology 36: 114–130, 2021.

Erika-Florence M, Leech R, Hampshire A. A functional network perspective on response inhibition and attentional control. Nat Commun 5: 4073, 2014.

Farina D, Negro F, Gizzi L, Falla D. Low-frequency oscillations of the neural drive to the muscle are increased with experimental muscle pain. J Neurophysiol 107: 958–965, 2012.

Feeney DF, Mani D, Enoka RM. Variability in common synaptic input to motor neurons modulates both force steadiness and pegboard time in young and older adults. J Physiology 596: 3793–3806, 2018.

Feingold J, Gibson DJ, DePasquale B, Graybiel AM. Bursts of beta oscillation differentiate postperformance activity in the striatum and motor cortex of monkeys performing movement tasks. Proc National Acad Sci 112: 13687–13692, 2015.

Fiori F, Chiappini E, Candidi M, Romei V, Borgomaneri S, Avenanti A. Long-latency interhemispheric interactions between motor-related areas and the primary motor cortex: a dual site TMS study. Sci Rep-uk 7: 14936, 2017.

Fitzgerald PB, Fountain S, Daskalakis ZJ. A comprehensive review of the effects of rTMS on motor cortical excitability and inhibition. Clin Neurophysiol 117: 2584–2596, 2006.

Frank MJ, Samanta J, Moustafa AA, Sherman SJ. Hold Your Horses: Impulsivity, Deep Brain Stimulation, and Medication in Parkinsonism. Science 318: 1309–1312, 2007.

Galganski ME, Fuglevand AJ, Enoka RM. Reduced control of motor output in a human hand muscle of elderly subjects during submaximal contractions. J Neurophysiol 69: 2108–2115, 1993.

Guan Y, Wessel JR. Two types of motor inhibition after action errors in humans. J Neurosci JN-RM-1191-22, 2022.

Hallett M, Chokroverty S. Magnetic Stimulation in Clinical Neurophysiology. 2nd ed. [Online]. American Journal of Neuroradiology 27: 1799, 2006 http://www.ajnr.org/content/27/8/1799.abstract.

Hamada M, Galea JM, Lazzaro VD, Mazzone P, Ziemann U, Rothwell JC. Two Distinct Interneuron Circuits in Human Motor Cortex Are Linked to Different Subsets of Physiological and Behavioral Plasticity. J Neurosci 34: 12837–12849, 2014.

Hannah R, Rothwell JC. Pulse Duration as Well as Current Direction Determines the Specificity of Transcranial Magnetic Stimulation of Motor Cortex during Contraction. Brain Stimul 10: 106–115, 2017.

Herz DM, Tan H, Brittain J-S, Fischer P, Cheeran B, Green AL, FitzGerald J, Aziz TZ, Ashkan K, Little S, Foltynie T, Limousin P, Zrinzo L, Bogacz R, Brown P. Distinct mechanisms mediate speed-accuracy adjustments in cortico-subthalamic networks. Elife 6: e21481, 2017.

Huster RJ, Enriquez-Geppert S, Lavallee CF, Falkenstein M, Herrmann CS. Electroencephalography of response inhibition tasks: Functional networks and cognitive contributions. Int J Psychophysiol 87: 217– 233, 2013.

Hyngstrom AS, Kuhnen HR, Kirking KM, Hunter SK. Functional implications of impaired control of submaximal hip flexion following stroke. Muscle Nerve 49: 225–232, 2014.

Ilmoniemi RJ, Hernandez-Pavon JC, Mäkelä NN, Metsomaa J, Mutanen TP, Stenroos M, Sarvas J. Dealing with Artifacts in TMS-Evoked EEG. 2015 37th Annu Int Conf Ieee Eng Medicine Biology Soc Embc 2015: 230–233, 2015.

Jahanshahi M, Rothwell JC. Inhibitory dysfunction contributes to some of the motor and non-motor symptoms of movement disorders and psychiatric disorders. Philosophical Transactions Royal Soc B Biological Sci 372: 20160198, 2017.

Jana S, Hannah R, Muralidharan V, Aron AR. Temporal cascade of frontal, motor and muscle processes underlying human action-stopping. Elife 9: e50371, 2020.

Jong RD, Coles MGH, Logan GD, Gratton G. In Search of the Point of No Return: The Control of Response Processes. J Exp Psychology Hum Percept Perform 16: 164–182, 1990.

Kok A, Ramautar JR, Ruiter MBD, Band GPH, Ridderinkhof KR. ERP components associated with successful and unsuccessful stopping in a stop-signal task. Psychophysiology 41: 9–20, 2004.

Kouzaki M, Shinohara M. Steadiness in plantar flexor muscles and its relation to postural sway in young and elderly adults. Muscle Nerve 42: 78–87, 2010.

Lazzaro VD, Rothwell JC. Corticospinal activity evoked and modulated by non-invasive stimulation of the intact human motor cortex. J Physiology 592: 4115–4128, 2014.

Leocani L, Cohen LG, Wassermann EM, Ikoma K, Hallett M. Human corticospinal excitability evaluated with transcranial magnetic stimulation during different reaction time paradigms. Brain 123: 1161–1173, 2000.

Leventhal DK, Gage GJ, Schmidt R, Pettibone JR, Case AC, Berke JD. Basal Ganglia Beta Oscillations Accompany Cue Utilization. Neuron 73: 523–536, 2012.

Logan GD, Cowan WB, Davis KA. On the ability to inhibit simple and choice reaction time responses: A model and a method. J Exp Psychology Hum Percept Perform 10: 276–291, 1984.

Majid DSA, Cai W, George JS, Verbruggen F, Aron AR. Transcranial Magnetic Stimulation Reveals Dissociable Mechanisms for Global Versus Selective Corticomotor Suppression Underlying the Stopping of Action. Cereb Cortex 22: 363–371, 2012.

Mani D, Almuklass AM, Hamilton LD, Vieira TM, Botter A, Enoka RM. Motor unit activity, force steadiness, and perceived fatigability are correlated with mobility in older adults. J Neurophysiol 120: 1988–1997, 2018.

Matzke D, Verbruggen F, Logan GD. Stevens’ Handbook of Experimental Psychology and Cognitive Neuroscience. 1–45, 2018.

McFarland DJ, Miner LA, Vaughan TM, Wolpaw JR. Mu and Beta Rhythm Topographies During Motor Imagery and Actual Movements. Brain Topogr 12: 177–186, 2000.

Miocinovic S, Hemptinne C de, Chen W, Isbaine F, Willie JT, Ostrem JL, Starr PA. Cortical Potentials Evoked by Subthalamic Stimulation Demonstrate a Short Latency Hyperdirect Pathway in Humans. J Neurosci 38: 9129–9141, 2018.

Nambu A, Tokuno H, Takada M. Functional significance of the cortico–subthalamo–pallidal ‘hyperdirect’ pathway. Neurosci Res 43: 111–117, 2002.

Novembre G, Pawar V, Bufacchi R, Kilintari M, Srinivasan M, Rothwell J, Haggard P, Iannetti G. Saliency detection as a reactive process: unexpected sensory events evoke cortico-muscular coupling. J Neurosci 38: 2474–17, 2018.

Novembre G, Pawar VM, Kilintari M, Bufacchi RJ, Guo Y, Rothwell JC, Iannetti GD. The effect of salient stimuli on neural oscillations, isometric force, and their coupling. Neuroimage 198: 221–230, 2019.

Pfurtscheller G, Graimann B, Huggins JE, Levine SP, Schuh LA. Spatiotemporal patterns of beta desynchronization and gamma synchronization in corticographic data during self-paced movement. Clin Neurophysiol 114: 1226–1236, 2003.

Picazio S, Veniero D, Ponzo V, Caltagirone C, Gross J, Thut G, Koch G. Prefrontal Control over Motor Cortex Cycles at Beta Frequency during Movement Inhibition. Curr Biol 24: 2940–2945, 2014.

Raud L, Huster RJ, Ivry RB, Labruna L, Messel MS, Greenhouse I. A Single Mechanism for Global and Selective Response Inhibition under the Influence of Motor Preparation. J Neurosci 40: 7921–7935, 2020.

Rogasch NC, Sullivan C, Thomson RH, Rose NS, Bailey NW, Fitzgerald PB, Farzan F, Hernandez-Pavon JC. Analysing concurrent transcranial magnetic stimulation and electroencephalographic data: A review and introduction to the open-source TESA software. Neuroimage 147: 934–951, 2017.

Rossi S, Antal A, Bestmann S, Bikson M, Brewer C, Brockmöller J, Carpenter LL, Cincotta M, Chen R, Daskalakis JD, Lazzaro VD, Fox MD, George MS, Gilbert D, Kimiskidis VK, Koch G, Ilmoniemi RJ, Lefaucheur JP, Leocani L, Lisanby SH, Miniussi C, Padberg F, Pascual-Leone A, Paulus W, Peterchev AV, Quartarone A, Rotenberg A, Rothwell J, Rossini PM, Santarnecchi E, Shafi MM, Siebner HR, Ugawa Y, Wassermann EM, Zangen A, Ziemann U, Hallett M, 2020 T basis of this article began with a CS from the IW on “Present Future of TMS: Safety, Ethical Guidelines”, Siena, October 17-20, 2018, updating through April. Safety and recommendations for TMS use in healthy subjects and patient populations, with updates on training, ethical and regulatory issues: Expert Guidelines. Clin Neurophysiol 132: 269–306, 2020.

Sherman MA, Lee S, Law R, Haegens S, Thorn CA, Hämäläinen MS, Moore CI, Jones SR. Neural mechanisms of transient neocortical beta rhythms: Converging evidence from humans, computational modeling, monkeys, and mice. Proc National Acad Sci 113: E4885–E4894, 2016.

Shin H, Law R, Tsutsui S, Moore CI, Jones SR. The rate of transient beta frequency events predicts behavior across tasks and species. Elife 6: e29086, 2017.

Siegert S, Ruiz MH, Brücke C, Huebl J, Schneider G-H, Ullsperger M, Kühn AA. Error signals in the subthalamic nucleus are related to post-error slowing in patients with Parkinson’s disease. Cortex 60: 103–120, 2014.

Skinner JW, Christou EA, Hass CJ. Lower Extremity Muscle Strength and Force Variability in Persons With Parkinson Disease. J Neurol Phys Ther 43: 56–62, 2019.

Smith M-C, Stinear CM. Transcranial magnetic stimulation (TMS) in stroke: Ready for clinical practice? J Clin Neurosci 31: 10–14, 2016.

Soh C, Hynd M, Rangel BO, Wessel JR. Adjustments to Proactive Motor Inhibition without Effector-Specific Foreknowledge Are Reflected in a Bilateral Upregulation of Sensorimotor β-Burst Rates. J Cognitive Neurosci 33: 784–798, 2021.

Swann N, Tandon N, Canolty R, Ellmore TM, McEvoy LK, Dreyer S, DiSano M, Aron AR. Intracranial EEG Reveals a Time- and Frequency-Specific Role for the Right Inferior Frontal Gyrus and Primary Motor Cortex in Stopping Initiated Responses. J Neurosci 29: 12675–12685, 2009.

Swann NC, Cai W, Conner CR, Pieters TA, Claffey MP, George JS, Aron AR, Tandon N. Roles for the pre-supplementary motor area and the right inferior frontal gyrus in stopping action: Electrophysiological responses and functional and structural connectivity. Neuroimage 59: 2860–2870, 2012.

Tatz JR, Soh C, Wessel JR. Common and unique inhibitory control signatures of action-stopping and attentional capture suggest that actions are stopped in two stages. J Neurosci JN-RM-1105-21, 2021.

Verbruggen F, Aron AR, Band GP, Beste C, Bissett PG, Brockett AT, Brown JW, Chamberlain SR, Chambers CD, Colonius H, Colzato LS, Corneil BD, Coxon JP, Dupuis A, Eagle DM, Garavan H, Greenhouse I, Heathcote A, Huster RJ, Jahfari S, Kenemans JL, Leunissen I, Li C-SR, Logan GD, Matzke D, Morein-Zamir S, Murthy A, Paré M, Poldrack RA, Ridderinkhof KR, Robbins TW, Roesch M, Rubia K, Schachar RJ, Schall JD, Stock A-K, Swann NC, Thakkar KN, Molen MW van der, Vermeylen L, Vink M, Wessel JR, Whelan R, Zandbelt BB, Boehler CN. A consensus guide to capturing the ability to inhibit actions and impulsive behaviors in the stop-signal task. Elife 8: e46323, 2019.

Volz LJ, Hamada M, Rothwell JC, Grefkes C. What Makes the Muscle Twitch: Motor System Connectivity and TMS-Induced Activity. Cereb Cortex 25: 2346–2353, 2015.

Wager TD, Sylvester C-YC, Lacey SC, Nee DE, Franklin M, Jonides J. Common and unique components of response inhibition revealed by fMRI. Neuroimage 27: 323–340, 2005.

Wagner J, Wessel JR, Ghahremani A, Aron AR. Establishing a Right Frontal Beta Signature for Stopping Action in Scalp EEG: Implications for Testing Inhibitory Control in Other Task Contexts. J Cognitive Neurosci 30: 107–118, 2018.

Wessel JR. β-Bursts Reveal the Trial-to-Trial Dynamics of Movement Initiation and Cancellation. J Neurosci 40: 411–423, 2020.

Wessel JR, Aron AR. Unexpected Events Induce Motor Slowing via a Brain Mechanism for Action-Stopping with Global Suppressive Effects. J Neurosci 33: 18481–18491, 2013.

Wessel JR, Aron AR. It’s not too late: The onset of the frontocentral P3 indexes successful response inhibition in the stop-signal paradigm. Psychophysiology 52: 472–480, 2015.

Wessel JR, Aron AR. On the Globality of Motor Suppression: Unexpected Events and Their Influence on Behavior and Cognition. Neuron 93: 259–280, 2017.

Wessel JR, Diesburg DA, Chalkley NH, Greenlee JDW. A causal role for the human subthalamic nucleus in non-selective cortico-motor inhibition. Curr Biol 32: 3785-3791.e3, 2022.

Wessel JR, Jenkinson N, Brittain J-S, Voets Shem, Aziz TZ, Aron AR. Surprise disrupts cognition via a fronto-basal ganglia suppressive mechanism. Nat Commun 7: 11195, 2016.

Wessel JR, Reynoso HS, Aron AR. Saccade suppression exerts global effects on the motor system. J Neurophysiol 110: 883–890, 2013.

Wessel JR, Waller DA, Greenlee JD. Non-selective inhibition of inappropriate motor-tendencies during response-conflict by a fronto-subthalamic mechanism. Elife 8: e42959, 2019.

Wilson JM, Thompson CK, McPherson LM, Zadikoff C, Heckman CJ, MacKinnon CD. Motor Unit Discharge Variability Is Increased in Mild-To-Moderate Parkinson’s Disease. Front Neurol 11: 477, 2020.

